# Deep Augmentation for Electrode Shift Compensation in Transient High-density sEMG: Towards Application in Neurorobotics

**DOI:** 10.1101/2022.02.25.481922

**Authors:** Tianyun Sun, Jacqueline Libby, JohnRoss Rizzo, S. Farokh Atashzar

## Abstract

Going beyond the traditional sparse multichannel peripheral human-machine interface that has been used widely in neurorobotics, high-density surface electromyography (HD-sEMG) has shown significant potential for decoding upper-limb motor control. We have recently proposed heterogeneous temporal dilation of LSTM in a deep neural network architecture for a large number of gestures (>60), securing spatial resolution and fast convergence. However, several fundamental questions remain unanswered. One problem targeted explicitly in this paper is the issue of “electrode shift,” which can happen specifically for high-density systems and during doffing and donning the sensor grid. Another real-world problem is the question of transient versus plateau classification, which connects to the temporal resolution of neural interfaces and seamless control. In this paper, for the first time, we implement gesture prediction on the transient phase of HD-sEMG data while robustifying the human-machine interface decoder to electrode shift. For this, we propose the concept of deep data augmentation for transient HD-sEMG. We show that without using the proposed augmentation, a slight shift of 10mm may drop the decoder’s performance to as low as 20%. Combining the proposed data augmentation with a 3D Convolutional Neural Network (CNN), we recovered the performance to 84.6% while securing a high spatiotemporal resolution, robustifying to the electrode shift, and getting closer to large-scale adoption by the end-users, enhancing resiliency.

## I. Introduction

In the last decade, thanks to significant technical and technological developments, neural interfaces used in neurorobotics have gone through major transformations, unleashing the potential for interfacing human cognition with machine intelligence. Focusing on peripheral interfacing in this paper, two major directions of research are (a) utilization of dense and flexible electronics and (b) implementation of deep biosignal processing, which can demystify the neural code of motor intention to be used for the control of neurorobots.

The field has gone through a rapid progression after the introduction of deep learning (DL), for example, [1]–[5]. Despite recent algorithmic progress, a major obstacle to real-world implementation is the need for extensive recalibration and the corresponding sensitivity of the trained model to various experimental conditions. Among the existing challenges is the topic of electrode shift, which is significantly more pronounced when signals are collected using miniaturized and densely-located electrodes. Electrode shift can be the result of doffing and donning the device, or it may happen due to skin stretches during intensive tasks. It is known that electrode shift would lead to degradation in model accuracy [6]–[8]. In order to address the issue of electrode displacement and misplacement, recalibration has been utilized in practice. However, it is a time-consuming process, complicated for non-expert users to conduct accurately, and cannot address several issues such as skin stretch. This has made the performance of such technologies limited and often increased the rejection rates of prosthetic systems. Thus, it can be summarized that there is an unmet need for the development of computational models and frameworks that minimize the need for recalibration and can generalize the performance over “possible electrode shift” configurations. The challenge of the electrode shift has been discussed in the literature for over a decade, initially on multi-channel electrodes sparsely placed directly on the muscles of interest. For example, the electrode shift issue has been discussed in the context of gesture and motion prediction using traditional ML models, where manual feature engineering was conducted to enhance the performance. In this regard, Young *et al*. [9] proposed optimization of the electrode configuration, achieving 78% accuracy on seven classes. In addition, there exists some literature that suggests fixing the problem by collecting a larger, more representative dataset.

Besides the concerns for generalizability and the limited number of gestures that were detected using classic algorithms, there are also concerns about the long data collection processes, which can be arduous for the end-user [10]–[13]. To avoid this, more recently, Ameri *et al*. [14] proposed the use of transfer learning for sparse multi-channel recording to minimize the amount of new data that is needed each time the user doffs/dons the device, allowing for a faster recalibration process. While the proposed transfer learning is an effective later-stage processing step, the compounded performance would be limited by the functionality of the initial phase of processing. It should also be noted that in most of these examples since the channels are treated independently, the spatial relationship between the channels is not fully exploited. This can be another limiting factor in considering electrode shift, which is inherently a spatial problem.

Fig. 1 shows an example of high-density sEMG (HD-sEMG), which has significantly boosted the information rate captured from one muscle. HD-sEMG enables decomposition of the collected dense signal space into spike train activities deriving motor units in the muscles [15]. More recently, HD-sEMG has been used as a direct input into deep neural network models to decode the motor intention of the user with high spatiotemporal resolution [1]. In [1], we have recently proposed a novel deep recurrent neural network, one of the very first deep neural networks on transient-phase high-density EMG data, that can map the dynamic phase of high-density recording captured using 128 channels from the upper limb (64 flexors and 64 extensors), into a prediction of over 60 classes of gestures. Utilizing the new concept of temporal dilation of LSTM, the proposed algorithm was able to make the convergence of the network 20 times faster when compared with conventional networks while providing high accuracy and sensitivity. In addition, our previous work showed that the transient phase of sEMG (which is classically discarded for ML-based processing) indeed has enough information to be used for augmenting the temporal resolution of human-machine interfaces. We also showed that the high-density information context significantly boosted the power of such machines so that the model could achieve over 80% performance for 65 gestures.

**Fig. 1:**
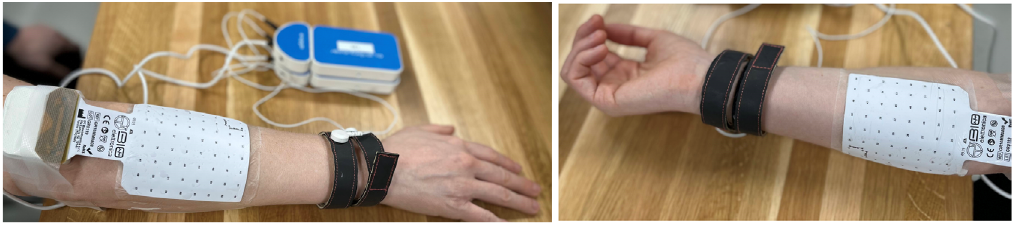
Two wireless OT Bioelettronica high-density sEMG 8×8 grids attached to the extensors (top) and flexors (bottom) of the forearm for gesture prediction.

The problem of electrode shift is significantly more pronounced for high-density recording of surface electromyography due to the size of the electrodes and since some existing computational models may depend on the corresponding contextual meaning of electrode locations. This issue is discussed in [16], which sheds light on the effect of electrode shift for decoding muscle synergies. The authors of [16] showed that when electrodes are shifted, it would require a new model for synergy estimation, which emphasizes the sensitivity of the problem to small electrode shifts and the existing central question regarding the generalizability of such models to various configurations.

In addition to the literature regarding the effect of electrode shift on decomposition and synergy, there exists some work showing the degradation of HD-sEMG-based decoders in the presence of small shifts of the sensor grid when using ML for gesture classification. Some have used classic ML algorithms and manual feature engineering. In this regard, Stango *et al*. [17] used spatial correlation features fed into Support Vector Machines, achieving 80% accuracy on 9 classes. Pan *et al*. [18] used Common Spacial Pattern features with LDA classifiers, achieving 80% accuracy on 11 classes. Also, He & Zhu [19] used Fourier domain features, achieving 85% accuracy on 11 classes. Also, Lv *et al*. [20] proposes memory-free autoencoder design of multilayer perceptron, achieving 90% accuracy for 10 classes. In the mentioned examples, the cost to address electrode shift was the lower number of degrees of freedom, limiting the versatility of the human-robot interface. [17]– [21]. However, these simplifications defeat the very purpose of neural interfaces, i.e., dexterity in predicting human motor intention.

In addition to the above, it can be mentioned that the existing literature mainly focuses on the plateau phase of the sEMG signals. However, a hand gesture involves a dynamic transient phase, which represents the dynamical recruitment of motor units in the first part of the task and contains critical temporal information. However, in the literature, this part of data is often discarded due to the complexity of modeling. Thus, it can be mentioned that gesture prediction from the transient phase is an under-explored area of research. Addressing this would allow for detecting the motor intent at the very beginning of the muscle contraction and when the signals are not stabilized. Such an approach would enable faster response and better real-time control. Thus, detecting abrupt transient phases of actions and translating that into motor intention is of high importance.

In this paper, for the first time, we address the problem of electrode shift for decoding the transient HD-sEMG while securing a large number of gesture classes. We showed that the performance of DL models could drop significantly (from %80 to %20) by a slight electrode shift. The specific augmentation proposed in this paper targets this issue while not requiring extra data collection. Utilizing a 3D CNN architecture, the proposed approach can robustly predict the gestures based on the transient HD-sEMG.

## II. Material and Methods

### A. Data Acquisition

In this paper, we used a high-density HD-sEMG database, available online [22], to allow benchmarking the proposed technique. The dataset contains 65 isometric hand gestures with different degrees of freedom (DoFs), including six 1-DoF finger and wrist gestures, forty-one 2-DoF compound gestures of fingers and wrist, and eight multi-DoF gestures of grasping, pointing, and pinching. Fig. 2 shows examples of these gestures: at rest, 1-Dof little finger bend, 1-Dof ring finger bend, and the 2-Dof combination of these two finger-bends. The HD-sEMG signals were recorded using a Quattrocento (OT Bioelettronica) biomedical amplifier system through two pads of 8×8 electrode grids (for a total of 128 channels) with a 10mm inter-electrode distance. As an example, a high-density sEMG system is depicted in Fig. 1 to highlight the electrode locations. The signals are collected at a sampling rate of 2048 Hz. This system allows for collecting dense biosignal space with information in both space and time. The two electrode grids are placed on both sides of the forearm, namely the dorsal (outer extensor muscles) and the volar (inner flexor muscles) of the upper forearm. In order to reduce common-mode noise, the recording was performed in a *differential manner*, where the signal of channel *i* in the final output signal is the difference between electrode *i* + 1 and *i* in the original collection. Twenty healthy adults, fourteen males and six females, with an age range between 25 and 57 years old, provided the data for collection. In this paper, we only used data from 19 subjects because data from subject 5 is unavailable. Each subject was asked to perform the gestures for five repetitions before switching to the next one. Each of those repetitions would last for 5 seconds, followed by an equal-duration interval of rest. In this way, the muscles get less influenced by fatigue and therefore provide more consistent data.

**Fig. 2:**
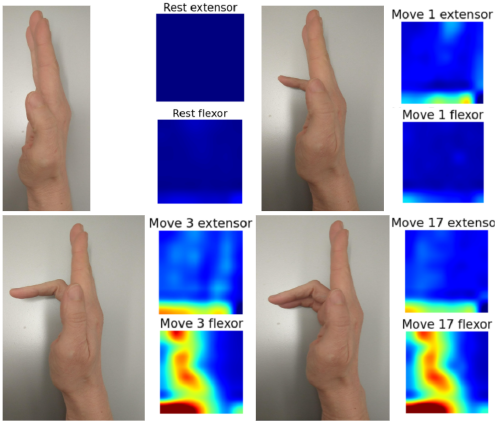
Examples of four gestures. (a) At rest; (b) Gesture 1: little finger bend; (c) Gesture 3: ring finger bend; (d) 2-Dof, Gesture 17: combining little finger and ring finger bend.

Some examples of muscle-activity heatmaps are shown in Fig. 3 for the best-performing subject to make the readers familiar with the heatmap representation of HD-sEMG. In this figure, due to the space constraints, we only show 16 out of the 65 gestures from the extensors and flexors grids, thus, a total of 32 heatmaps.

**Fig. 3:**
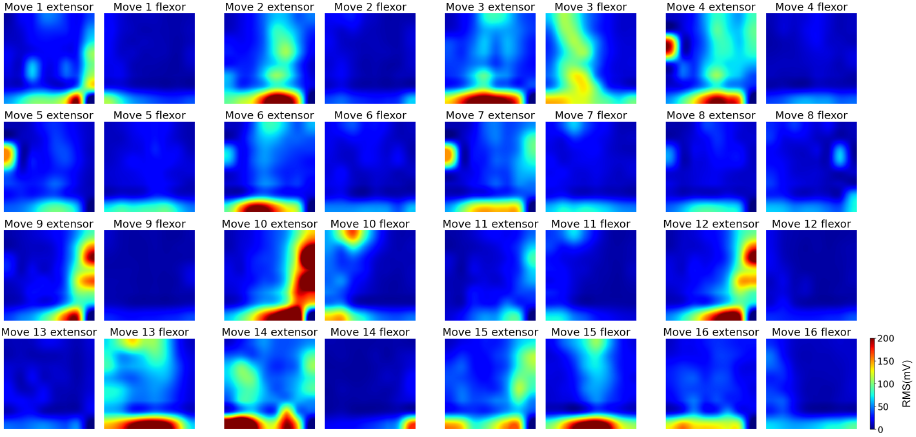
32 muscle-activity heatmaps associated with 16 1-DoF movements from the best-performing subject (#17). Each gesture has two heatmaps (forearm extensor and flexor). Each heatmap is an 8×8 grid, consisting 64 electrodes.

Forces measured at each finger and the wrist are included in the dataset to provide the labeling mechanism for cutting the transient and plateau phases of the data. The plateau phase is once the gesture has reached its final steady state, and the transient phase is the movement of the from rest to the begining of the steady state, as can be seen in Fig. 5.

### B. Data Preprocessing

As mentioned before, in this paper, we compare the performance of the proposed model on the transient phase with that of the model on the plateau phase. In this paper, we define these phases by averaging the force signal of each gesture (see Fig. 5). From this, we chose the first 0.5 seconds of data as the most dynamic portion of the sEMG time series, which encodes the transient phase. Force signals are shown in Fig. 5 for the same gestures shown in Fig. 2 (Movement 1: little finger bend, Movement 3: ring finger bend, and Movement 17: little finger bend plus ring finger bend.) The 0.5-second transient phase demarcation is shown with a dashed line. Force indices 0-8 denote measurements on the index finger, middle finger, ring finger, little finger, thumb finger flexion/extension, thumb finger abduction/adduction, wrist-flexion/extension, wrist-pronation/supination, and wrist-radial/ulnar, respectively.

In this paper, in order to normalize the signal space of HD-sEMG, we propose to use the *µ*-law transformation after applying a Min-Max normalization on the signals. *µ*-law transformation enhances the discriminability among information channels and therefore helps the model capture more information.

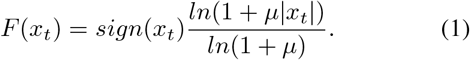

In (1), *x*_*t*_ denotes each data scalar and *µ* is a hyperparameter and selected to be 2048. Following the real-time implementation standards in myoelectric control, [23]–[27], we choose to conduct our experiments using a sliding window size of 200ms with a step size of 10ms. On each timestamp, one data point is generated, with the dimension of (sampling rate×window size)×8×8×2, where 8×8 is the size of each of the two grids. To help understand, Fig. 4 visualizes the data collection and sliding window process.

**Fig. 4:**
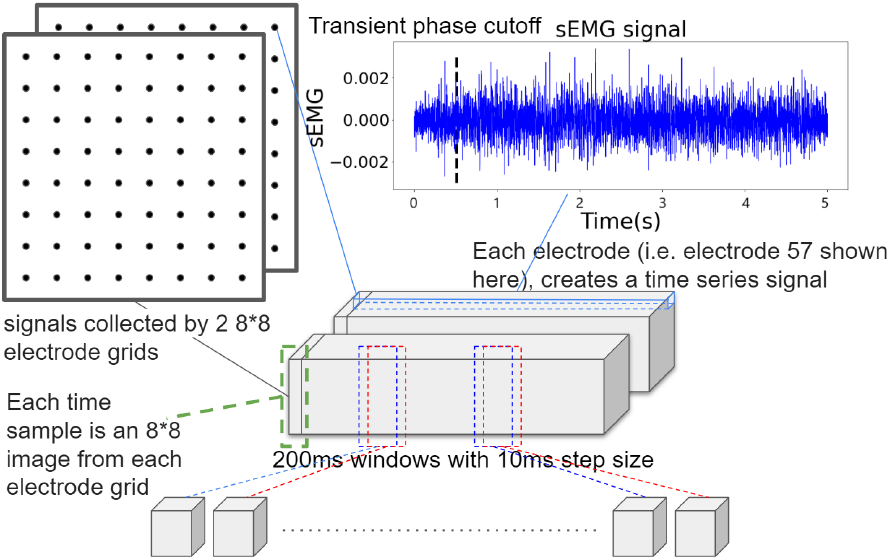
Data acquisition and sliding window process.

**Fig. 5:**
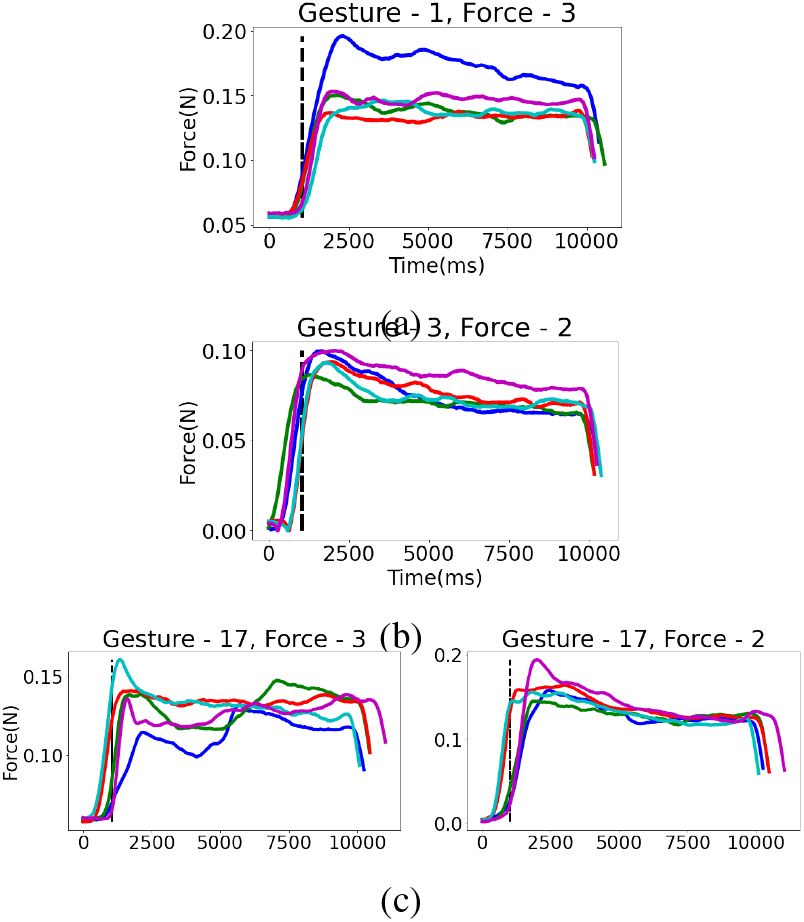
The corresponding forces of the same three gestures shown in Fig. 2, on each repetition. The dashed lines indicate the end (0.5 seconds) of transient phases. Line colors denote five different repetitions. (a) Force by little finger flexing; (b) Force by ring finger flexing; (c) Force by little finger and ring finger flexing.

### C. Data Augmentation

In order to augment the input space without requiring to collect a large data, instead of using the full set of 8×8 (which is the complete neurophysiological window for each electrode grid), we use subsets of 6×6 (named as the *window of observation*). This would leave out two extra electrodes which can be leveraged for data augmentation. In other words, the task will be to predict the gesture based on the window of observation, which may shift to right/left/top/down (or a dual combination) resembling electrode shift and augmenting the input space, which will force the neural network to learn the common underlying patterns of information which can be decoded in all shifted input space. In terms of the numbers, the aforementioned techniques would result in multiple choices of 6×6 observations window of observation within the 8×8 neurophysiological activation window. This data augmentation process is visually show in Fig. 6. Thus, two alternative spatial shifts can be considered, one includes only one-step shifting, and one includes two-step shifting. In the experiments where we use the one-step shifts, our dataset is multiplied by a factor of 5 (one ‘standard’ position of no shift + four ‘one-step’ shifts including shift to the right, left, top and down). In the experiments where we use both the one-step and two-step shifts, our dataset is multiplied by a factor of 9 (one ‘standard’ position + four ‘one-step’ shifts + four ‘two-step’ shifts).

**Fig. 6:**
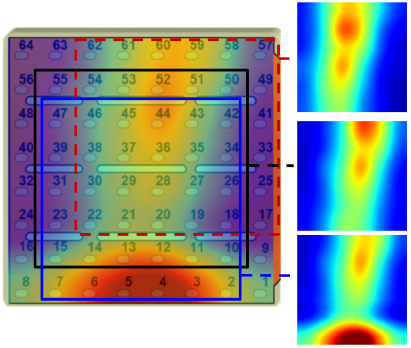
The heatmap with the 8×8 electrode grids. The black box shows the ‘standard’ position, the blue box shows a shift ‘one-step away’, and the dashed red line shows a shift ‘two-step away’. The corresponding heatmap of the boxes are shown on the right.

## III. Model Structure

Motivated by the literature on deep video processing, we considered the 2D signal at each timestamp as a frame of a neurophysiological video capturing the 2D spatial distributed muscle activation over time. In addition to 3D CNN we also tested the performance of 2D CNN deep neural networks, and a hybrid solution (when adding recurrent layers for their potential in modeling temporal dynamics). This is to conduct a comprehensive comparative analysis on the performance of the system taking into account complexity and performance.

### A. CNN-RNN Hybrid Models

Our 2D CNN-RNN hybrid model is depicted in Fig. 7a. The concept is to assign spatial decoding to CNN and temporal modeling to RNN. The CNN section is composed of a series of Conv2D layers with 2×2 kernel size and Relu activation function for nonlinearity, each with 8, 16, 32, 64 and 128 filters to gradually parse the images to a vector of 128 channels. It parses the (number of samples)×6×6×2-sized input signal to a shape of (number of samples)×72. Afterwards, it is fed into a 4-layer LSTM network with 128 hidden parameters and 400 LSTM nodes on each layer. Finally, the output of the LSTM layers is fed into 3 fully-connected layers with 65 nodes in the last layer, corresponding to the number of classes for the classification task. We also train a 1D CNN-RNN hybrid model. For this, instead of taking the 6×6 signal through a 2D CNN, the signals are reshaped into a 36×1 vector and then considered 1D convolution.

**Fig. 7:**
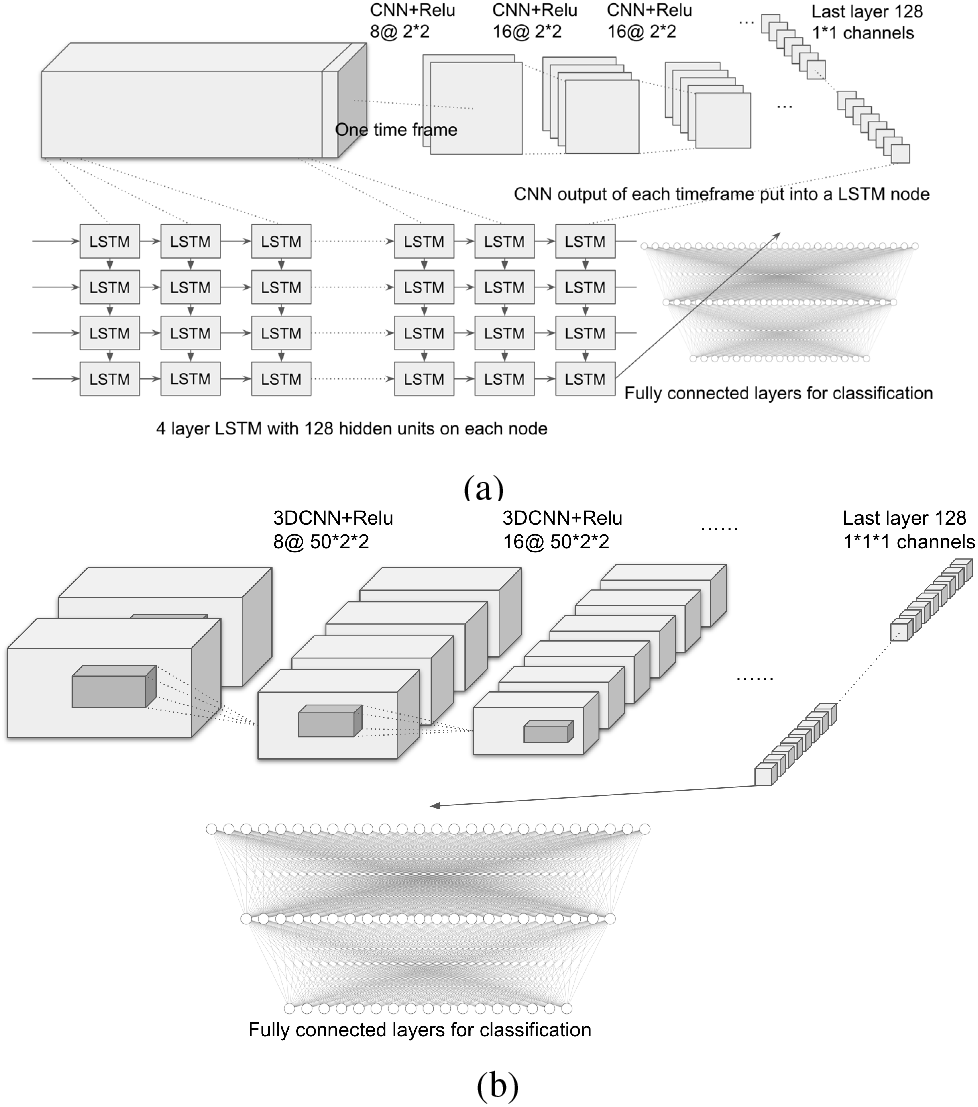
(a) Hybrid model using a combination of 2D CNN and multilayer LSTM; (b) Model using 3D CNN

### B. 3D CNN Model

In the case of 3D CNN models, as shown in Fig. 7b, we directly take the whole input data through several Conv3D layers to let the models decompose spatial and temporal information at the same time. This is critical to consider spatial and temporal interactions between various segments of the signal. In detail, the structure consists of 4 layers of (100, 2, 2) kernel size, each with 8, 16, 32 and 64 filters, followed by 1 layer of (4, 2, 2) kernel and 128 filters. Similar to the 2D case, the output was flattened and fed into three fully connected layers for classification.

## IV. Experiments and Results

### A. Experiment Models

In this part,the accuracy of the three aforementioned CNN models are compared using data with ‘one-step away’ augmentation. The experiment was conducted using bothe transient phase and stable phase data. In both cases, the 3D CNN structure consistently resulted in best performance. As the next step, the 3D CNN-based model was treined using both ‘1 step away’ and ‘2 steps away’ augmentations. In addition, the performance of a small experiment we call ‘non-aug’ is represented, where the best performing model was trained in the ‘standard’ position and tested in one of the ‘1 step away’ shifts to simulate the case of misplacement. The experimental models and the corresponding model ID’s are listed in table I. The model ID’s act as a reference for a later box plot. The median model accuracy across all experiment subjects of every experiment is listed in table II.

**TABLE I:** Model ID

|  | Conv1d | Conv2d | Conv3d | Conv3d 2step |
| --- | --- | --- | --- | --- |
| Transient | 1 | 2 | 3 | 4 |
| Stable | 5 | 6 | 7 | 8 |

**TABLE II:** Median model accuracy

|  | Non-Aug | Conv1D | Conv2D | Conv3D | Conv3D 2step |
| --- | --- | --- | --- | --- | --- |
| Transient | 18.54% | 69.01% | 67.89% | 78.50% | **80.22%** |
| Stable | 20.03% | 73.61% | 72.63% | **84.60%** | 84.45% |

### B. Results and Statistical Analysis

To evaluate the performances, k-fold (k=5) cross validation was utilized. For this, one of the five repetitions was held out for testing and the rest was used for training. Fig. 8 shows the box plots of the classification accuracy of the aforementioned models using either the transient phase signal or the stable phase signal. In the box plot, the model ID corresponds to what is listed in table I, the black horizontal lines indicate the medians, and the green triangles indicate the means. In order to evaluate the significance of effect from the proposed augmentation method, statistical analysis is performed on all the aforementioned models across all 19 subjects. Model 1 did not pass the Shapiro normality analysis. Thus, we performed Wilcoxon’s test for analyizing the significance. Bonferroni correction was applied to the observed p-values. The corrected p-value ranges are denoted with markers at the top of Fig 8. The marker symbols are defined as follows: (a) The ns marker (for not significant) represents 0.05 to 1; (b) * represents 0.01 to 0.05; (c) ** represents 0.001 to 0.01; (d) *** represents 0.0001 to 0.001; and (e) **** represents smaller than 0.0001.

**Fig. 8:**
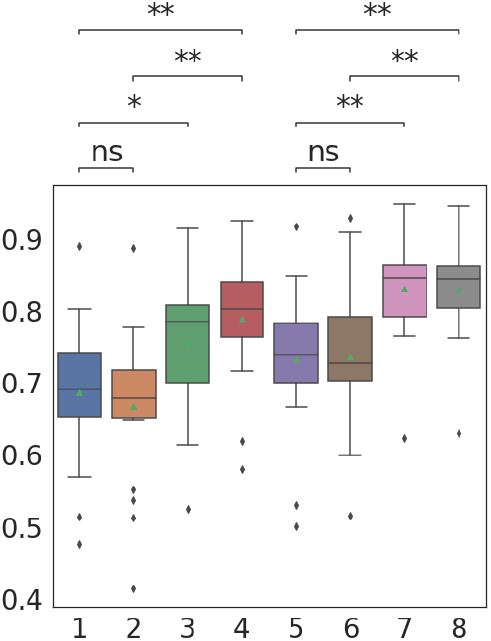
Classification accuracy of the experiment models.

As can be seen in the results, the performance gradually increases going from using 1D to 2D and to 3D Convolutional architectures. Conv1D and Conv2D did not show significant differences under both cases of transient and stable signals. The Conv3D model achieved the highest performance and significant gain compared to Conv1D and 2D. This can be due to the fact that Conv3D structure utilizes the full spectrum of spatial and temporal content of the signal. This can be particularly important when handling electrode shift for the dynamic phase since the spatial component is critical for handling the electrode shift, while the temporal is critical during the dynamic phase. We also observe that by adding data augmentations that are 2-steps away from the ‘standard position,’ the performance gained slight improvement.

## V. Conclusion

In this paper, we have implemented a data augmentation strategy to make the performance of HD-sEMG-based neural interfaces robust to electrode shift, focusing on the transient phase of the signal, which would also enhance the agility of the system. Using 128 HD-sEMG channels with the proposed data augmentation, a 3D CNN classifier was trained, which achieved 84.6% accuracy on the transient phase in the presence of synthetic electrode shift. The results showed that without such augmentation, the performance of the neural interface could significantly drop to as low as 20%.

